## Supplementary Figures, Tables and Videos for "Metallo-Supramolecular Branched Polymer Protects Particles from Air-water Interface in Single-Particle Cryo-Electron Microscopy"

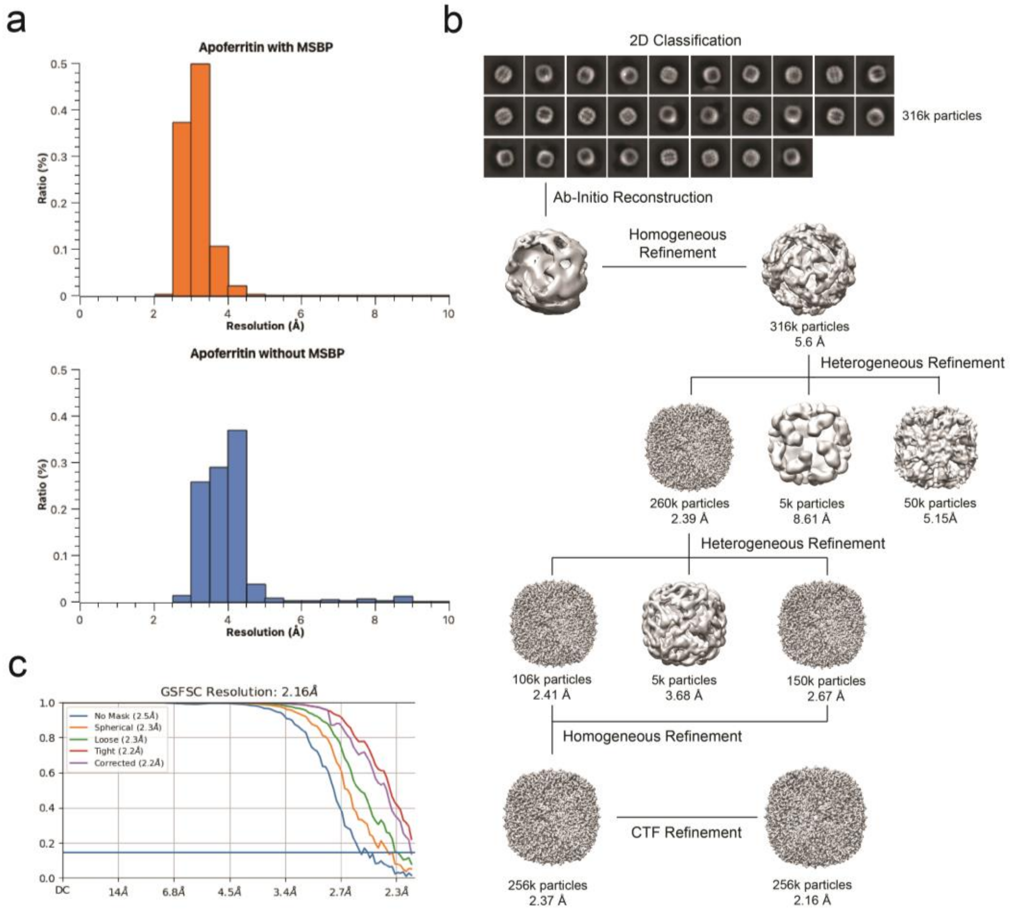


**Supplementary Fig.1 Cryo-EM data processing of apoferritin with MSBP.** **(a)** Statistics of estimated resolution of micrographs for apoferritin with or without MSBP. **(b)** Data processing workflow with number of particles and the reconstruction resolution indicated at every step. **(c)** FSC curves of the 3D reconstructions is also shown.

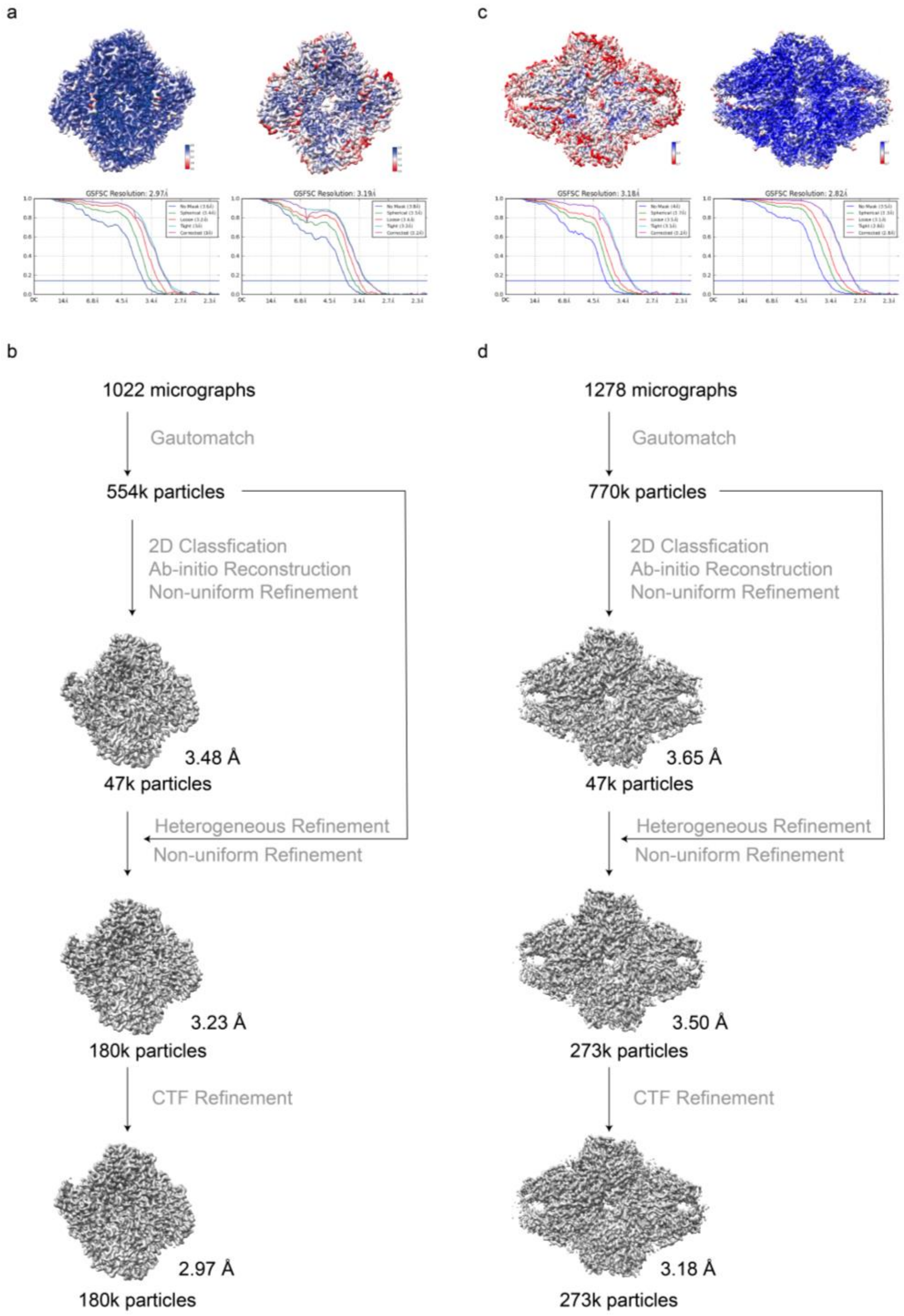


**Supplementary Fig.2 Cryo-EM data processing of catalase and β-galactosidase. (a)** Local resolution cryo-EM map and corresponding FSC curves of catalase with (left) or without (right) MSBP. **(b)** Data processing workflow of catalase with MSBP. Number of particles and the reconstruction resolution are indicated at every step. **(c)** Local resolution cryo-EM map and corresponding FSC curves of β-galactosidase with (left) or without (right) MSBP. **(d)** Data processing workflow of β-galactosidase with MSBP. Number of particles and the reconstruction resolution are indicated at every step.

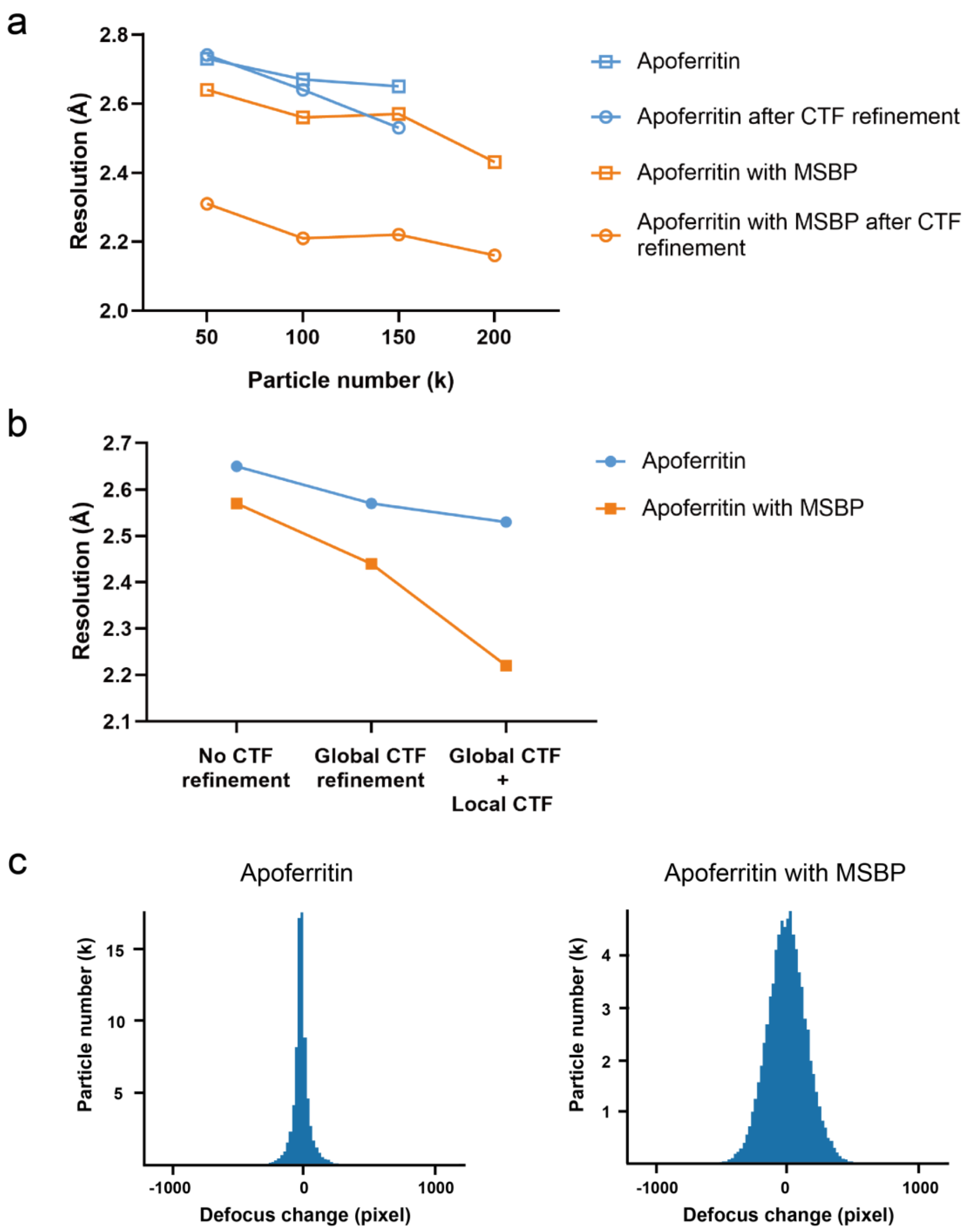


**Supplementary Fig.3 Systematically comparison of resolution for apoferritin with or without MSBP. (a)** The resolution corresponding to density map reconstructed with 50k, 100k, 150k and 200k particles are shown. Apoferritin with MSBP before (orange square) and after (orange circle) CTF refinement, apoferritin without MSBP before (blue square) and after (blue circle) ctf refinement are plotted for comparison. **(b)** Comparison of the resolution improvement by global and local CTF refinement with 150k particles of apoferritin (blue) and apoferritin with MSBP (orange). **(c)** Particle distribution of defocus changes after local CTF refinement for apoferritin (left) or apoferritin with MSBP (right).

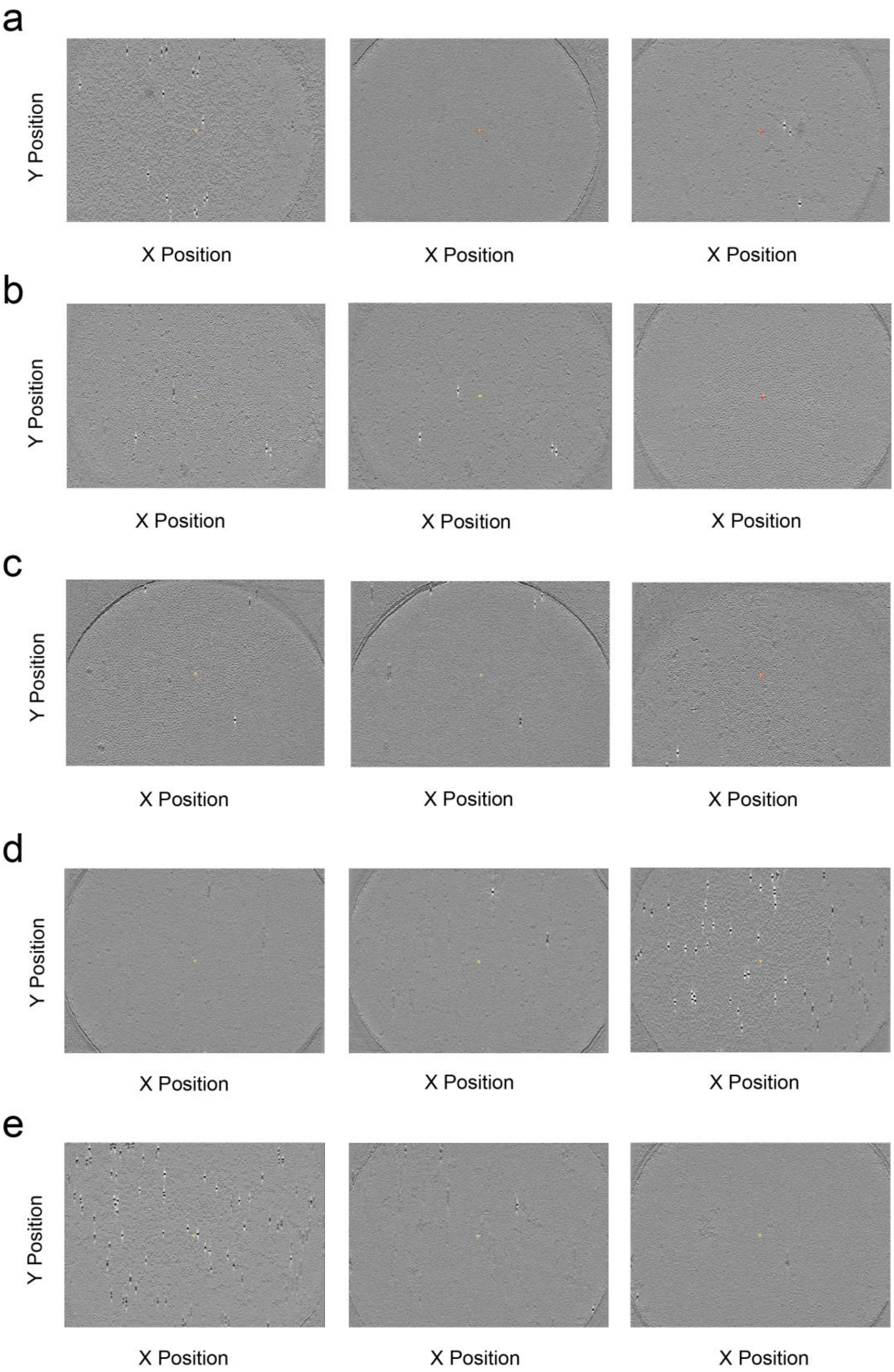


**Supplementary Fig.4 Particle distribution of apoferritin in three different layers of tomograms.** Comparison of images from upper air-water interface (left), inner (middle) and lower air-water interface (right) layers from tomograms for different datasets: apoferritin only (a), apoferritin with MSBP (b), MSBP only (c), apoferritin with PEG (d), apoferritin with $Pd(NO_3)_2$ (e).

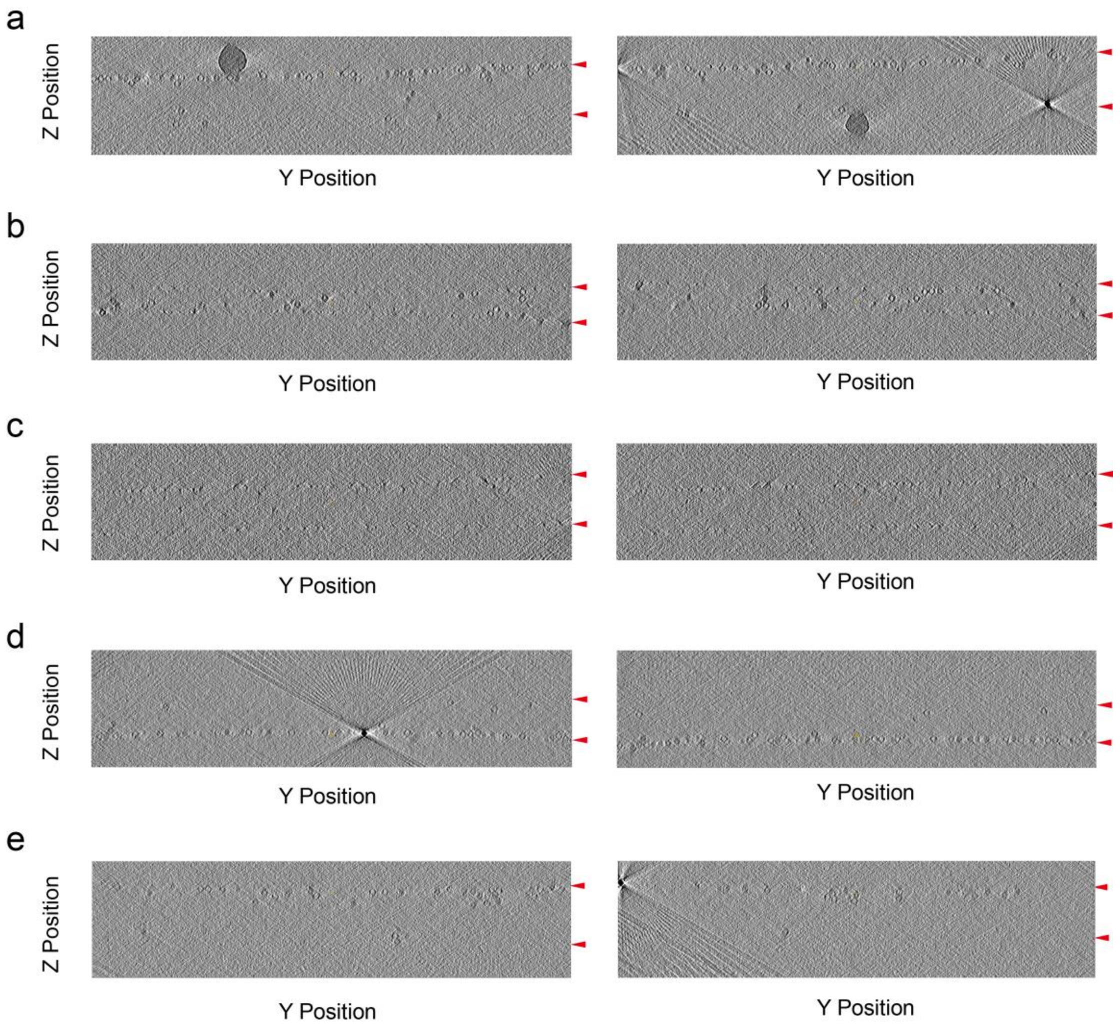


**Supplementary Fig.5 Side view (Y-Z) segmentations of tomograms.** Two different side view segmentations from tomograms for different datasets: apoferritin only (a), apoferritin with MSBP (b), MSBP only (c), apoferritin with PEG(d), apoferritin with $Pd(NO_3)_2$(e). Red triangles refer to the air-water interfaces.

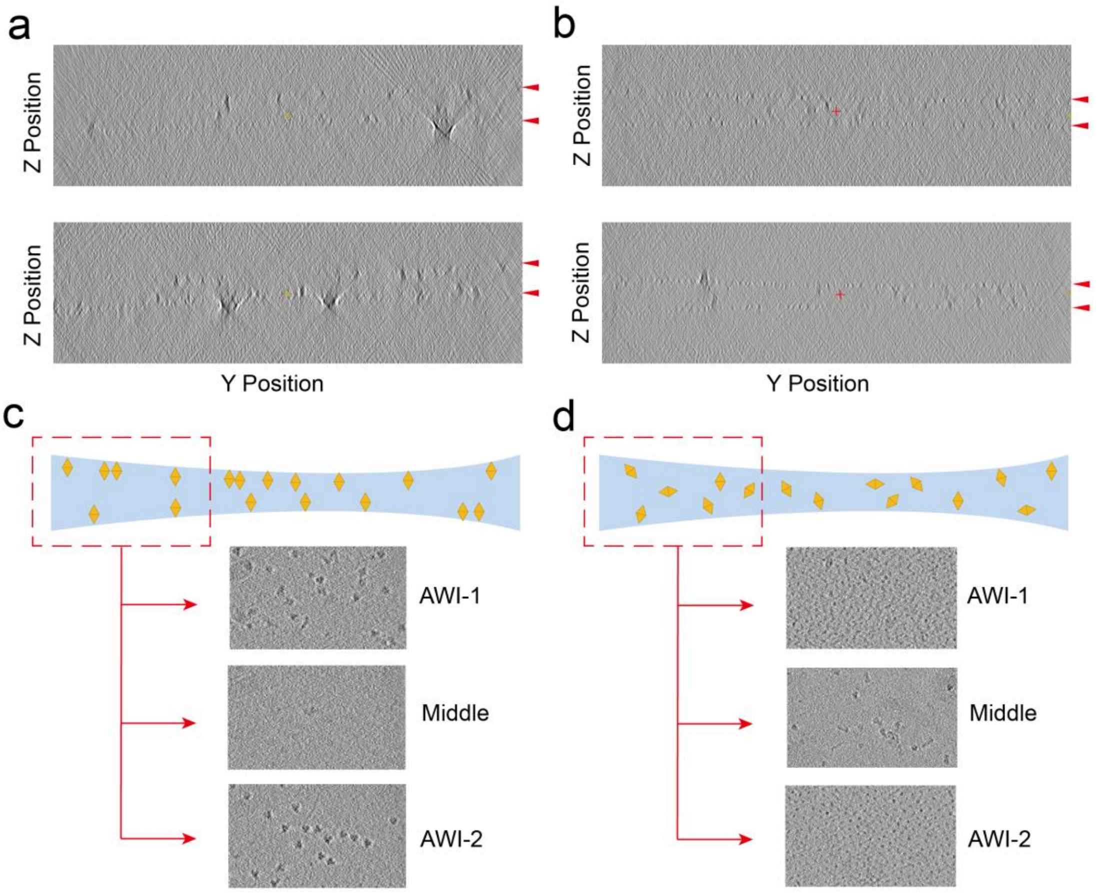


**Supplementary Fig.6 Particle distribution of HA trimer in vitreous ice.** Two different side view segmentations from tomograms of HA trimer without (a) or with (b) MSBP. (c) Schematic diagram shows particle distribution of HA trimer without MSBP in vitreous ice. Three layers along the Z axis of the tomogram indicated most of the particles are trapped in two AWIs, few particles are observed in middle of the vitreous ice. (d) Schematic diagram shows particle distribution of HA trimer with MSBP in vitreous ice. Three layers along the Z axis of the tomogram indicated particles are observed in middle of the vitreous ice.

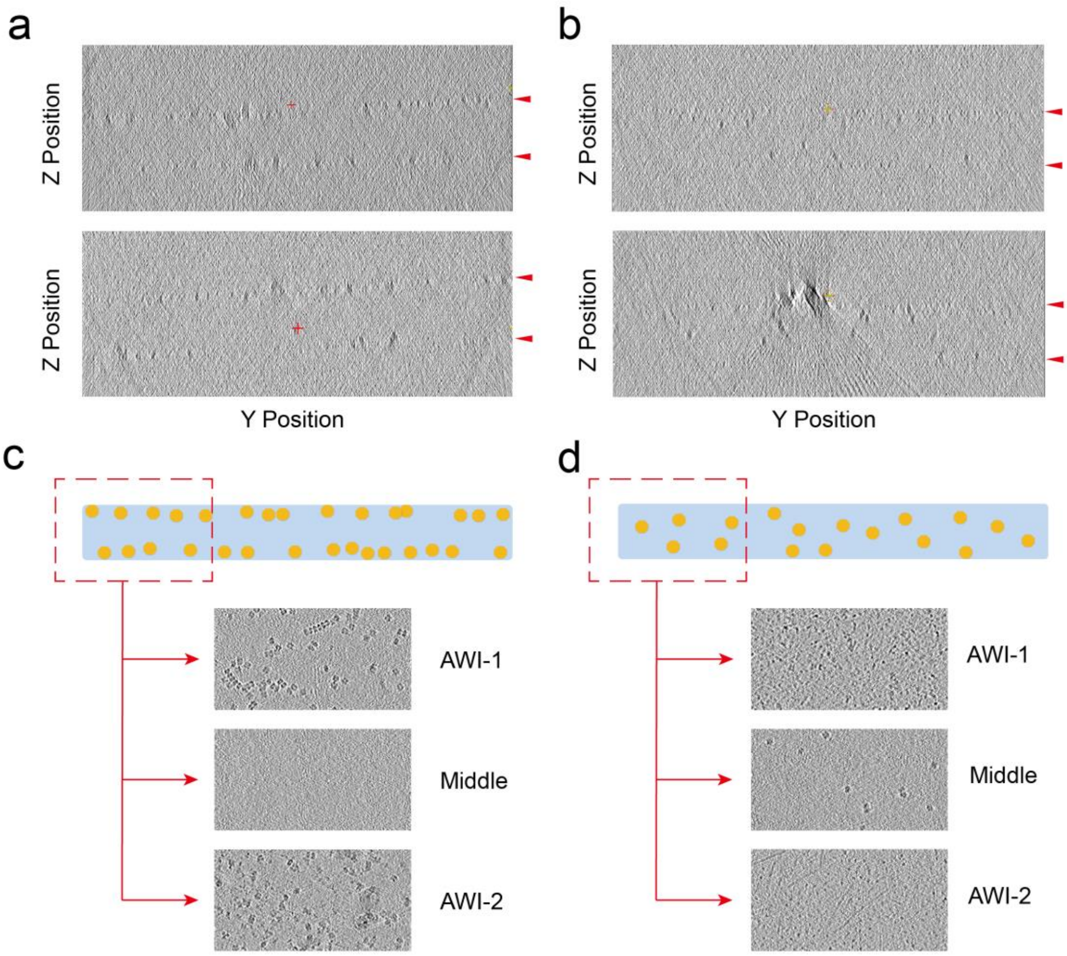


**Supplementary Fig.7 Particle distribution of catalase in vitreous ice.** Two different side view segmentations from tomograms of catalase without (a) or with (b) MSBP. (c) Schematic diagram shows particle distribution of catalase without MSBP in vitreous ice. Three layers along the Z axis of the tomogram indicated most of the particles are trapped in two AWIs, few particles are observed in middle of the vitreous ice. (d) Schematic diagram shows particle distribution of catalase with MSBP in vitreous ice. Three layers along the Z axis of the tomogram indicated particles are observed in middle of the vitreous ice.

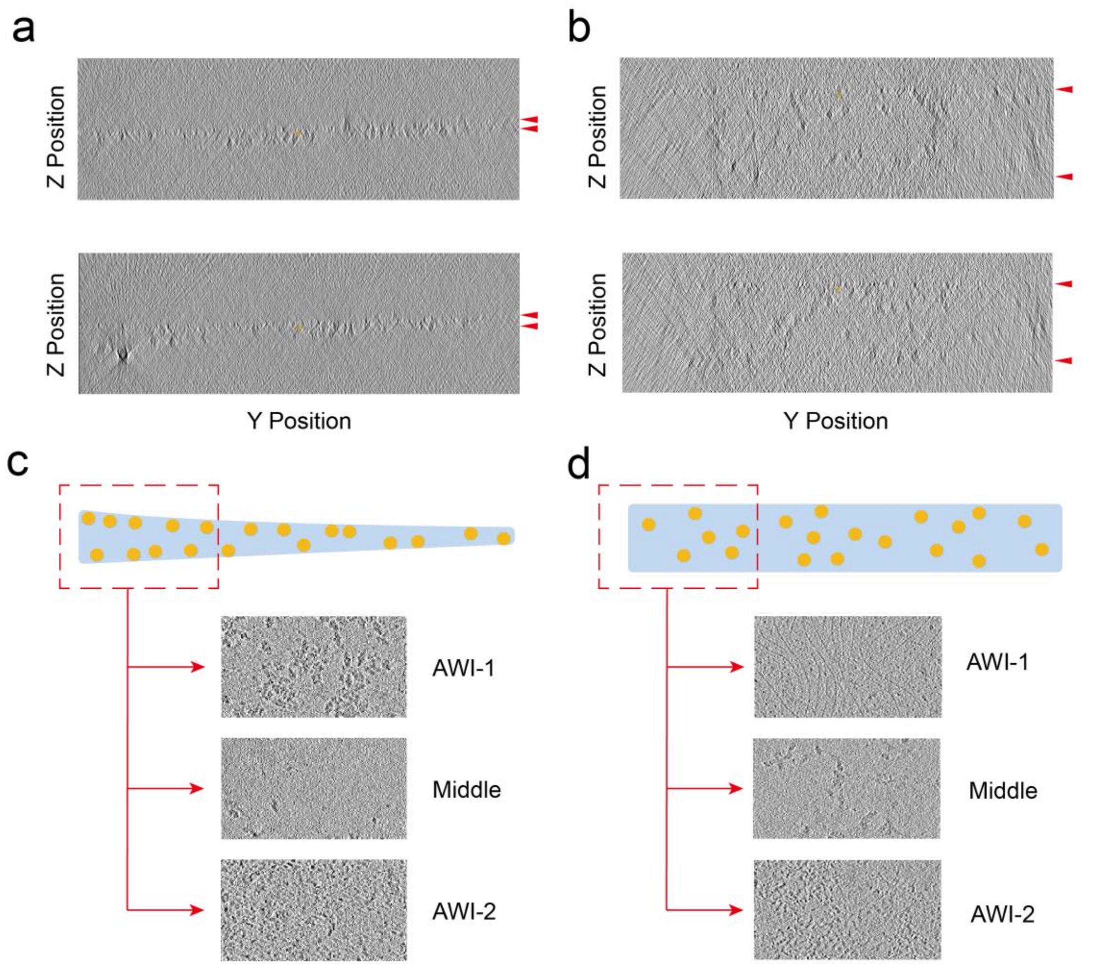


**Supplementary Fig.8 Particle distribution of β-galactosidase in vitreous ice.** Two different side view segmentations from tomograms of β-galactosidase without (a) or with (b) MSBP. (c) Schematic diagram shows particle distribution of β-galactosidase without MSBP in vitreous ice. Three layers along the Z axis of the tomogram indicated most of the particles are trapped in two AWIs, few particles are observed in middle of the vitreous ice. (d) Schematic diagram shows particle distribution of β-galactosidase with MSBP in vitreous ice. Three layers along the Z axis of the tomogram indicated particles are observed in middle of the vitreous ice.

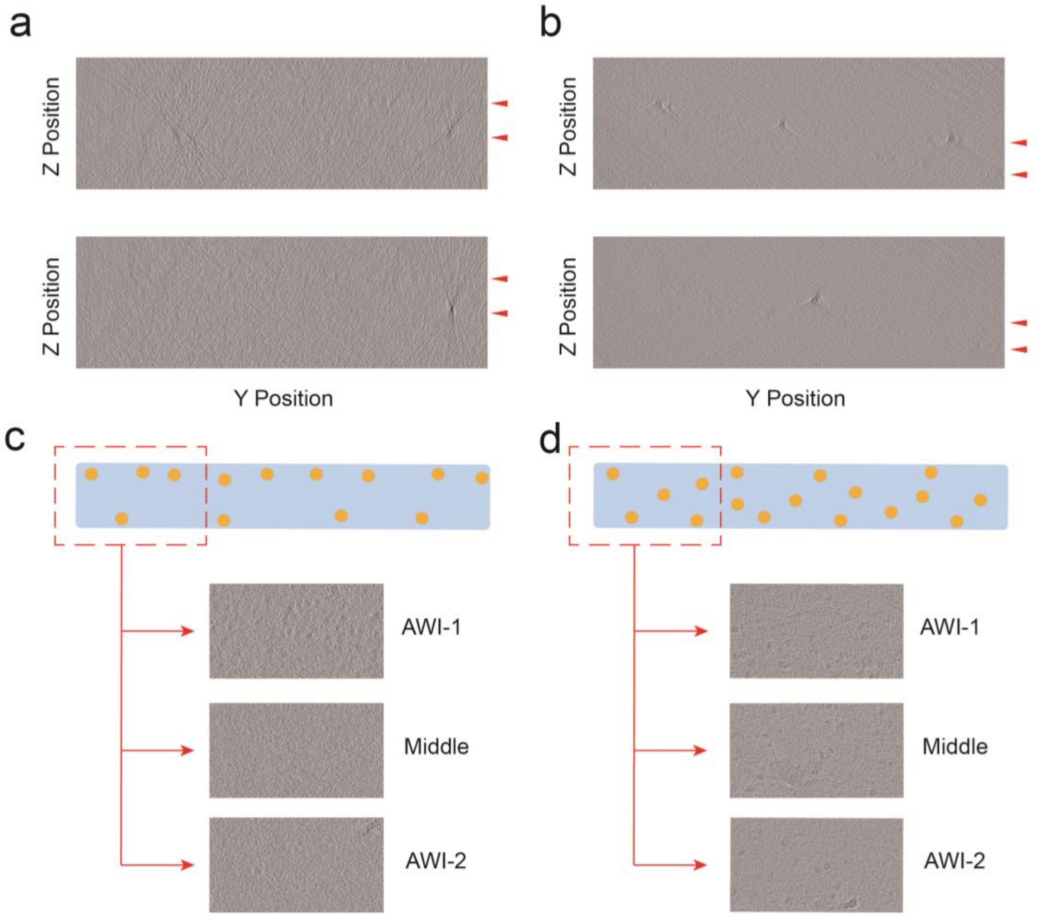


**Supplementary Fig.9 Particle distribution of IMP1 in vitreous ice.** Two different side view segmentations from tomograms of IMP1 without (a) or with (b) MSBP. (c) Schematic diagram shows particle distribution of IMP1 without MSBP in vitreous ice. Three layers along the Z axis of the tomogram indicated most of the particles are trapped in two AWIs, few particles are observed in middle of the vitreous ice. (d) Schematic diagram shows particle distribution of IMP1 with MSBP in vitreous ice. Three layers along the Z axis of the tomogram indicated particles are observed in middle of the vitreous ice.

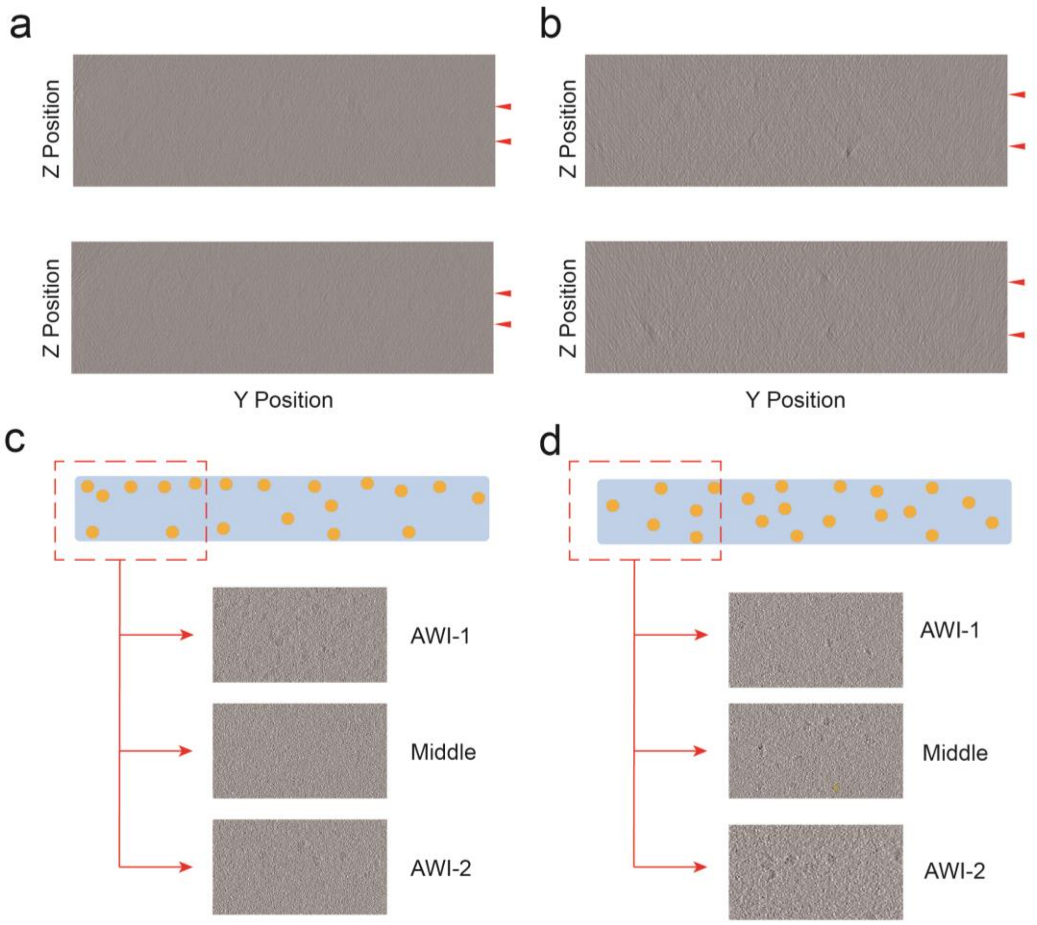


**Supplementary Fig.10 Particle distribution of IMP2 in vitreous ice.** Two different side view segmentations from tomograms of IMP2 without (a) or with (b) MSBP. (c) Schematic diagram shows particle distribution of IMP2 without MSBP in vitreous ice. Three layers along the Z axis of the tomogram indicated most of the particles are trapped in two AWIs, few particles are observed in middle of the vitreous ice. (d) Schematic diagram shows particle distribution of IMP2 with MSBP in vitreous ice. Three layers along the Z axis of the tomogram indicated particles are observed in middle of the vitreous ice.

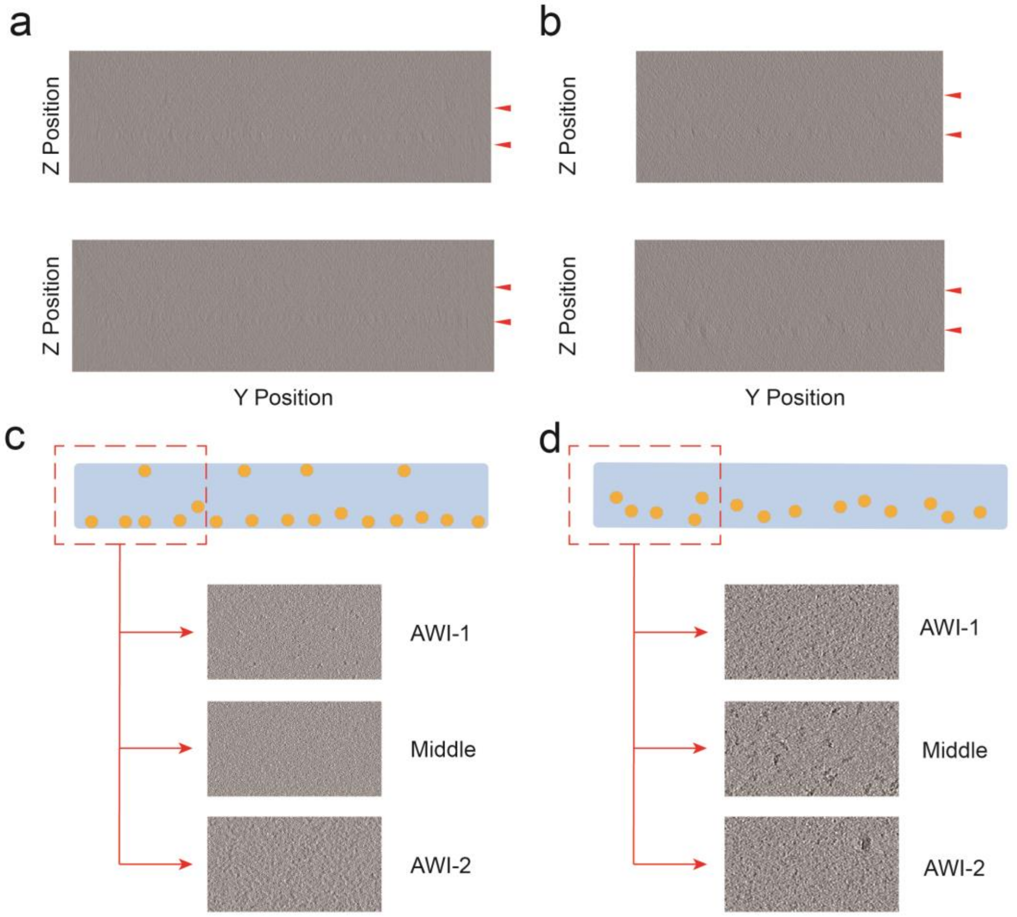


**Supplementary Fig.11 Particle distribution of CSW complex in vitreous ice.** Two different side view segmentations from tomograms of CSW complex without (a) or with (b) MSBP. (c) Schematic diagram shows particle distribution of CSW complex without MSBP in vitreous ice. Three layers along the Z axis of the tomogram indicated most of the particles are trapped in two AWIs, few particles are observed in middle of the vitreous ice. (d) Schematic diagram shows particle distribution of CSW complex with MSBP in vitreous ice. Three layers along the Z axis of the tomogram indicated particles are observed in middle of the vitreous ice.

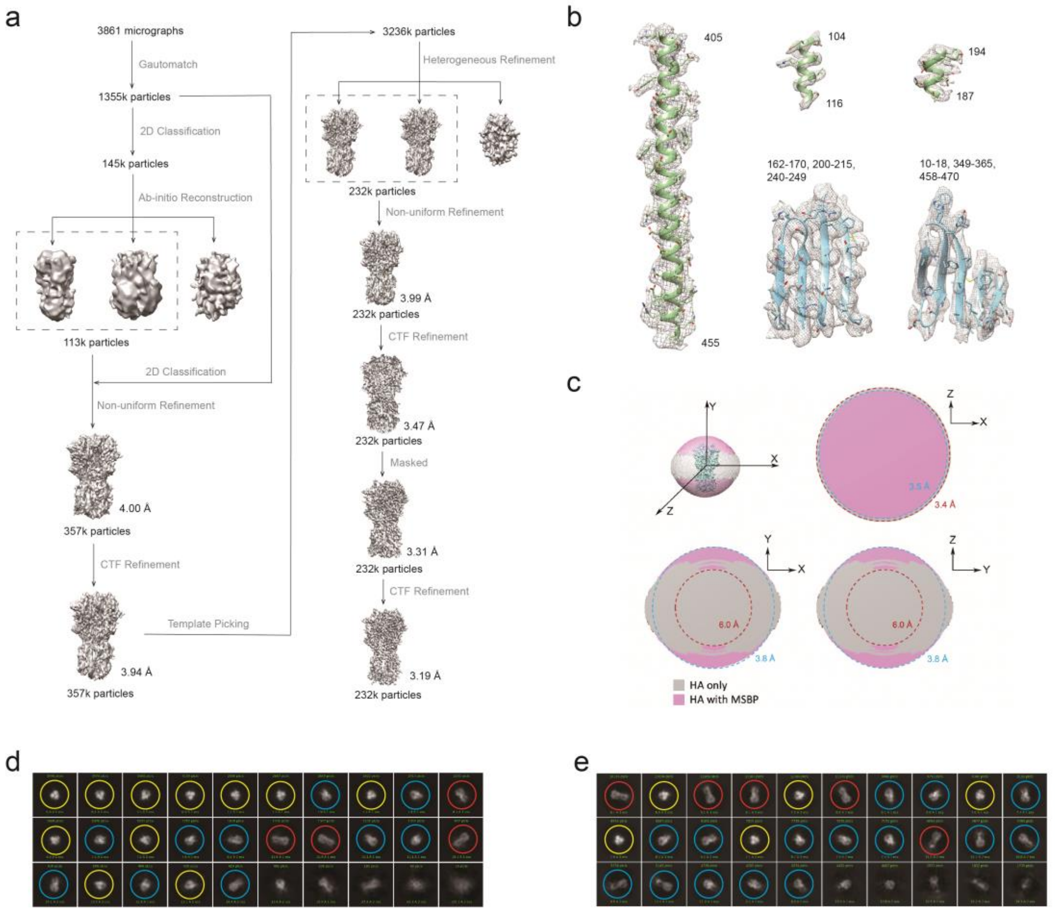


**Supplementary Fig.12 Cryo-EM data analysis of hemagglutinin trimer. (a)** Data processing workflow with number of particles and the reconstruction resolution indicated at every step. **(b)** Representative cryo-EM densities of HA with MSBP. **(c)** Calculated resolution from different views for HA trimer without (grey) or with (purple) MSBP. **(d-e)** 2D averages of HA trimer without (d) or with (e) MSBP. Top views (yellow circles), side views (red circles) and tilted views (blue circles) are indicated.

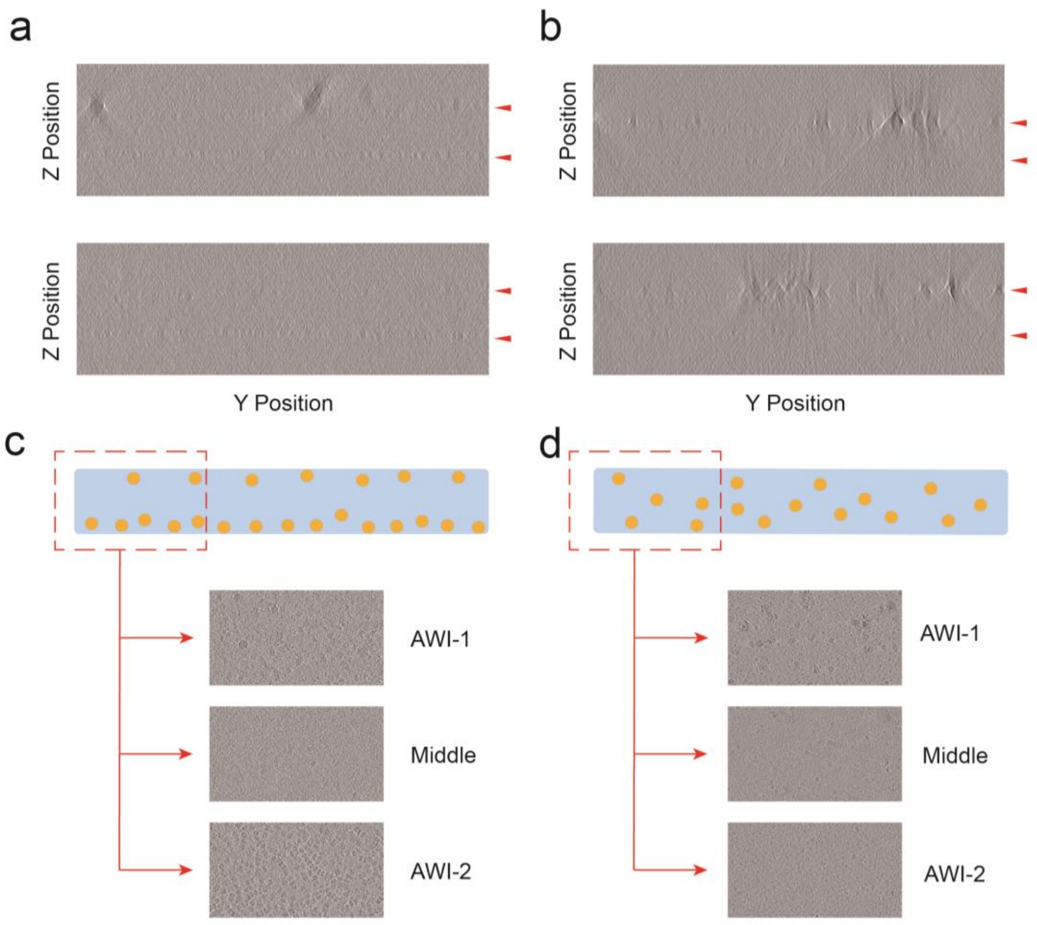


**Supplementary Fig.13 Particle distribution of catalase under high concentration of salt in vitreous ice.** Two different side view segmentations from tomograms of catalase prepared under high salt concentration (1M NaCl) without (a) or with (b) MSBP. (c) Schematic diagram shows particle distribution of catalase without MSBP in vitreous ice. Three layers along the Z axis of the tomogram indicated most of the particles are trapped in two AWIs, few particles are observed in middle of the vitreous ice. (d) Schematic diagram shows particle distribution of catalase with MSBP in vitreous ice. Three layers along the Z axis of the tomogram indicated particles are observed in middle of the vitreous ice.

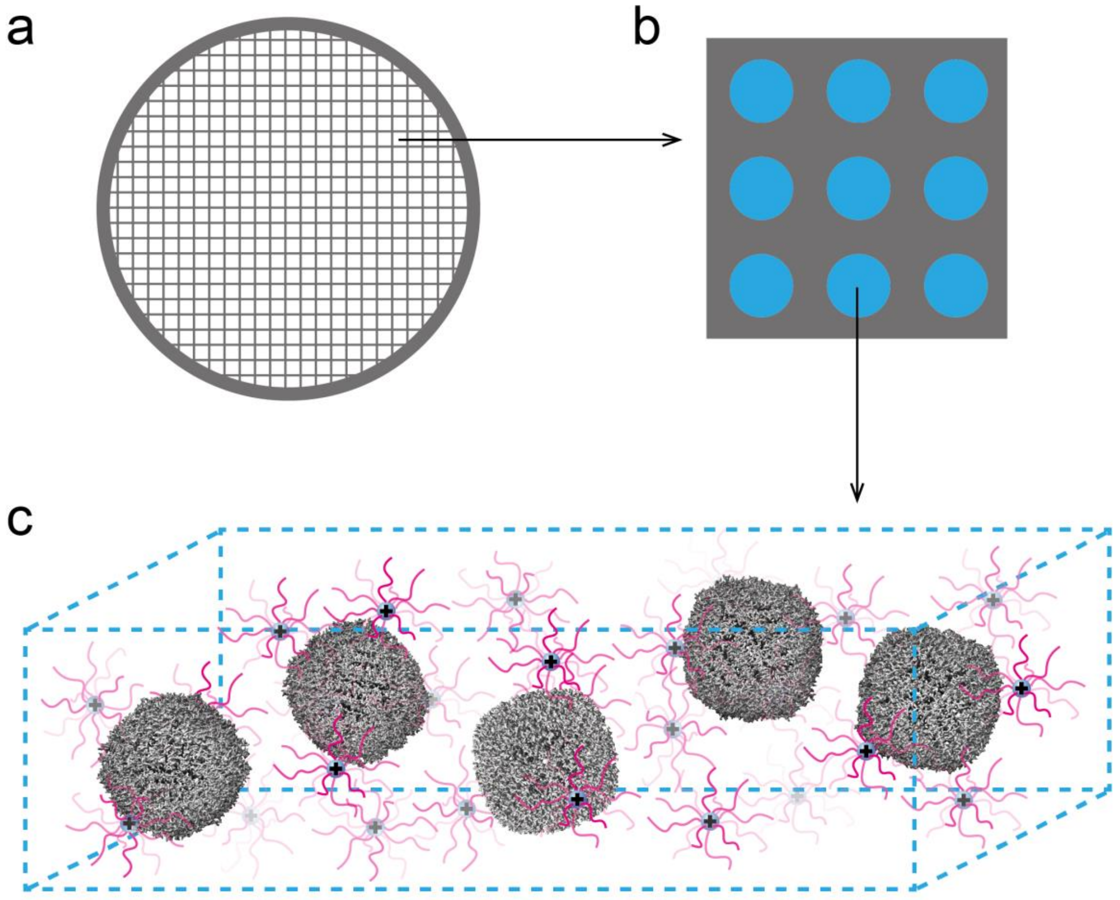


**Supplementary Fig.14 Proposed model of MSBP interacting with protein particles. (a)** A schematic of a typical holey carbon cryo-grids is shown. (**b**) A small region of cryo-grids with holes and carbon film shown as blude and grey, respectively. (**c**) Proposed protein particles in vitreous ice with MSBP applied. Protein particles are denoted as grey sphere, the MSBP nanoclusters are indicated as pink PEG polymers connected to positively charged part.

**Supplementary Table 1 Summary of Cryo-EM Data Collection and Processing.**

| **Protein** | Apoferritin | | Hemagglutinin | | Catalase | | β-Galactosidase | |
|---|---|---|---|---|---|---|---|---|
| MSBP | - | + | - | + | - | + | - | + |
| **Data collection** | | | | | | | | |
| EM equipment | Titan Krios | | | | | | | |
| Voltage (kV) | 300 | | | | | | | |
| Detector | Gatan K3 summit | | | | | | | |
| Magnification | 81000 x | | | | | | | |
| Pixel size (Å) | 1.06 | | | | | | | |
| Electron dose (e/Å²) | 57.04 | 57.04 | 51.75 | 50.9 | 50.58 | 50.58 | 50.58 | 50.63 |
| Defocus range (μm) | -1.3~-2.5 | -1.3~-2.5 | -1.3~-2.5 | -1.3~-2.5 | -1.0~-2.5 | -1.0~-2.5 | -1.0~-2.5 | -1.0~-2.5 |
| Collected movies | 2,090 | 1,587 | 576 | 3,861 | 1,053 | 1,022 | 1,317 | 1,278 |
| **Reconstruction** | | | | | | | | |
| Software | cryoSPARC v2.15.0, RELION 3.1 | | | | | | | |
| Final particles | 150,000 | 255,785 /150,000 | 127,885 | 231,931 | 180,161 | 180,161 | 273,058 | 273,058 |
| B-factors (Å²) | -127.5 | -87.9 /-88.4 | -141.5 | -149.5 | -127.5 | -129 | -112.6 | -131.6 |
| Map resolution (Å) | 2.53 | 2.16 /2.22 | 3.41 | 3.19 | 3.19 | 2.97 | 2.82 | 3.18 |
| EMDB code | | EMD-36313 | | EMD-36314 | | EMD-36315 | | EMD-36316 |

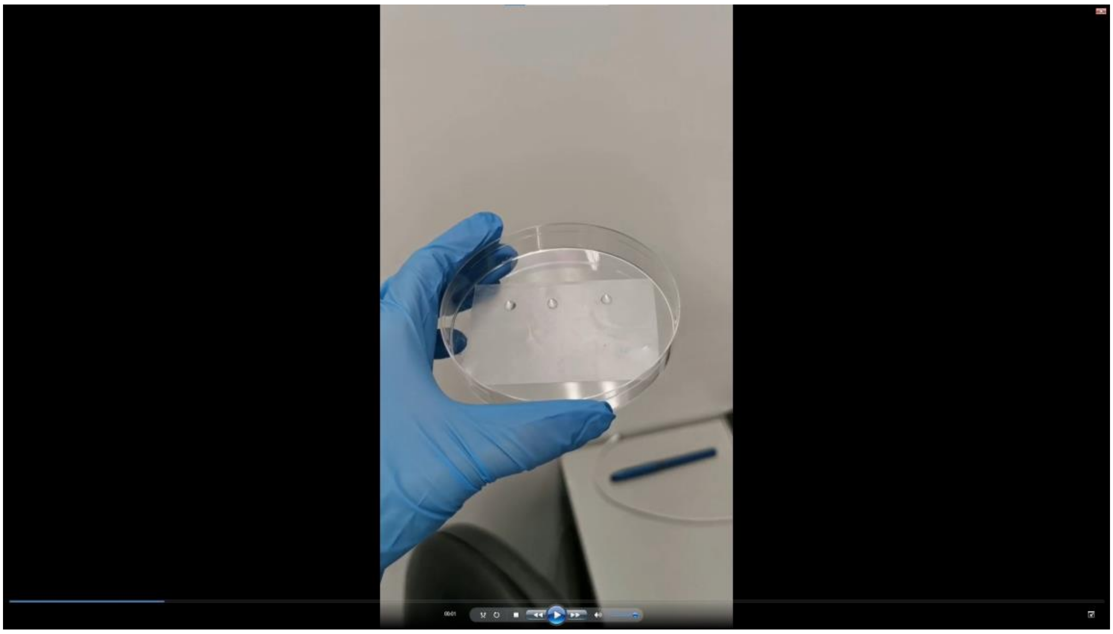

**Supplementary Video 1  Formation of MSBP in solution.** Three 20 μL drops are shown for comparison: 2 mg/mL PEG solution only (left), 0.24 mg/mL $Pd(NO_3)_2$ solution only (middle), mixture of 2 mg/mL PEG and 0.24 mg/mL $Pd(NO_3)_2$ solution (right).

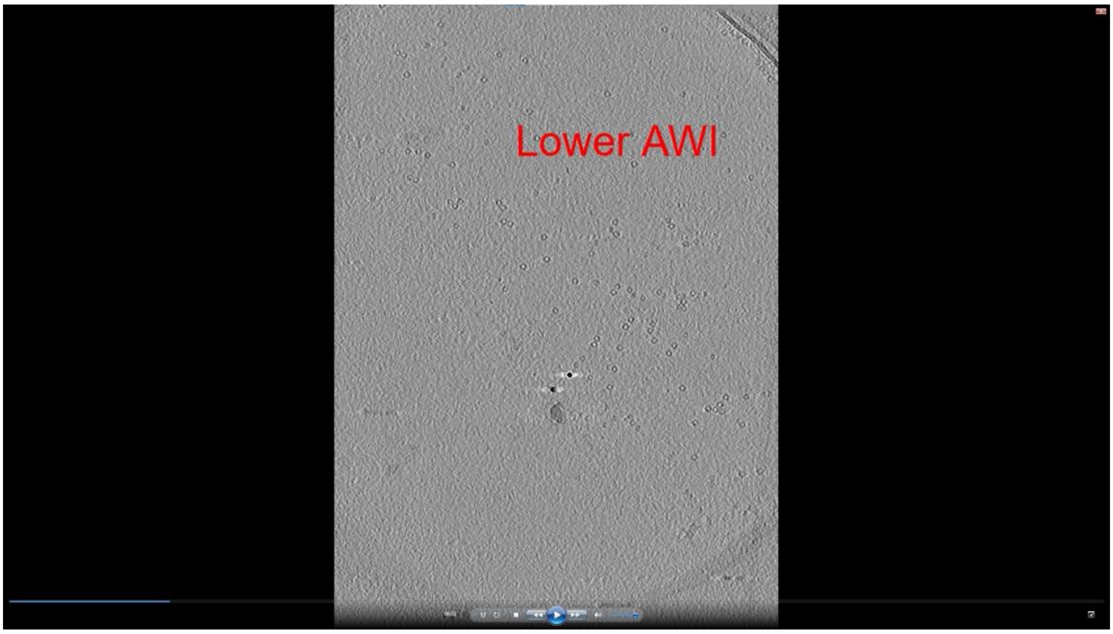


**Supplementary Video 2  Reconstructed tomogram of apoferritin without MSBP.** Lower and Upper AWIs are indicated.

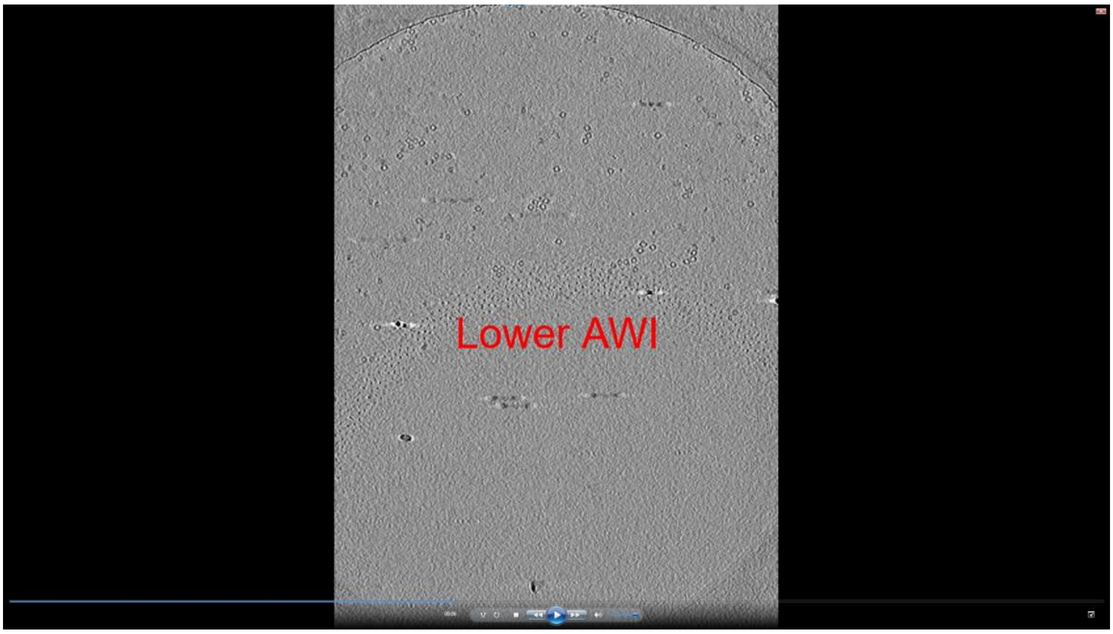


**Supplementary Video 3 Reconstructed tomogram of apoferritin with MSBP.** Lower and Upper AWIs are indicated.

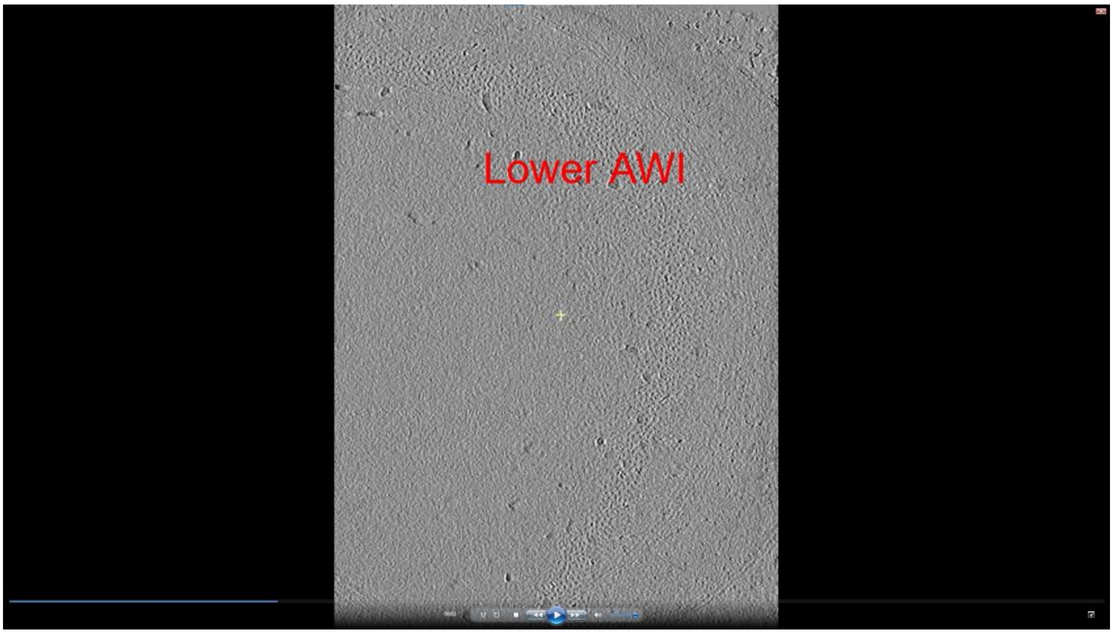


**Supplementary Video 4 Reconstructed tomogram of MSBP only.** Lower and Upper AWIs are indicated.

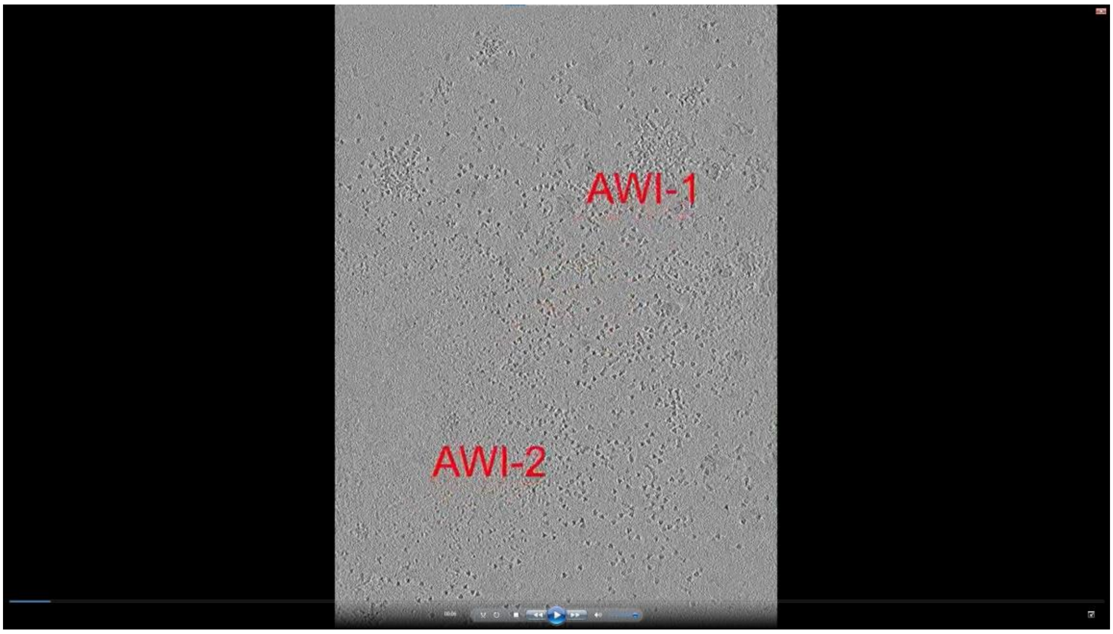


**Supplementary Video 5 Reconstructed tomogram of proteins used in this study**. Lower and Upper AWIs are indicated. The proteins shown in videos include HA (0:00:00-0:00:23), catalase (0:00:23-0:00:54), β-galactosidase (0:00:54-0:01:16), IMP1 (0:01:16-0:01:38) and IMP2 (0:01:38-0:02:25), and CSW (C9ORF72-SMCR8-WDR41) complex (0:02:25-0:03:05).
